# Modeling Cardiomyocyte Signaling and Metabolism Predicts Genotype to Phenotype Mechanisms in Hypertrophic Cardiomyopathy

**DOI:** 10.1101/2023.09.25.559356

**Authors:** A. Khalilimeybodi, Jeffrey J. Saucerman, P. Rangamani

## Abstract

Familial hypertrophic cardiomyopathy (HCM) is a significant precursor of heart failure and sudden cardiac death, primarily caused by mutations in sarcomeric and structural proteins. Despite the extensive research on the HCM genotype, the complex, context-specific nature of many signaling and metabolic pathways linking the HCM genotype to phenotype has hindered therapeutic advancements for patients. To address these challenges, here, we have developed a computational systems biology model of HCM at the cardiomyocyte level. Utilizing a stochastic logic-based ODE method, we integrate subcellular systems in cardiomyocytes that jointly modulate HCM genotype to phenotype, including cardiac signaling, metabolic, and gene regulatory networks, as well as posttranslational modifications linking these networks. After validating with experimental data on changes in activity of signaling species in HCM context and transcriptomes of two HCM mouse models (R403Q-αMyHC and R92W-TnT), the model predicts significant changes in cardiomyocyte metabolic functions such as ATP synthase deficiency and a transition from fatty acids to carbohydrate metabolism in HCM. The model indicated major shifts in glutamine-related metabolism and increased apoptosis after HCM-induced ATP synthase deficiency. Aligned with prior experimental studies, we predicted that the transcription factors STAT, SRF, GATA4, TP53, and FoxO are the key regulators of cardiomyocyte hypertrophy and apoptosis in HCM. Using the model, we identified shared (e.g., activation of PGC1*α* by AMPK, and FHL1 by titin) and context-specific mechanisms (e.g., regulation of Ca^2+^ sensitivity by titin in HCM patients) that could control genotype to phenotype transition in HCM across different species or mutations. We also predicted potential combination drug targets for HCM (e.g., mavacamten paired with ROS inhibitors) preventing or reversing HCM phenotype (i.e., hypertrophic growth, apoptosis, and metabolic remodeling) in cardiomyocytes. This study provides new insights into mechanisms linking genotype to phenotype in familial hypertrophic cardiomyopathy and offers a framework for assessing new treatments and exploring variations in HCM experimental models.

## 1 INTRODUCTION

Familial hypertrophic cardiomyopathy (HCM) is the most prevalent inherited cardiac disease affecting 1 in 500 up to 1 in 200 individuals and posing a potential risk to nearly 2 million individuals in the United States alone [1, 2]. HCM is a significant precursor of heart failure and the leading cause of sudden cardiac death in young adults [3, 4]. In HCM, gene mutations primarily at sarcomeric and structural proteins result in hypertrophic remodeling of cardiomyocytes which is characterized by the parallel addition of sarcomeres to increase left ventricular mass and wall thickness [5]. While this hypertrophic remodeling of cardiomyocytes can be an adaptive response to reduce cardiac wall stress, it is often maladaptive and an independent risk factor for cardiovascular morbidity and mortality [6]. Untreated HCM can potentially progress to heart failure, escalating morbidity and mortality [7]. Existing drug therapies often manage symptoms of hypertrophic cardiomyopathy rather than preventing or reversing cardiomyocyte growth and remodeling [8]. Despite significant progress in understanding the HCM genotype [9], the knowledge gap in the specific mechanisms linking genotype to phenotype in HCM hinders the development of new therapeutics. This gap potentially originates from 1) the significant complexity of genotype to phenotype mechanisms, regulated by a complex network of signaling and metabolic pathways [10, 11], and 2) the context dependency of these mechanisms influenced by variety in gene mutations, experimental models, and environmental factors [12, 13]. While past research has highlighted certain signaling and metabolic pathways that may link genotype and phenotype in HCM [12, 14–16], our understanding of how these pathways manifest across different contexts (i.e., mutations, and species), through a complex network of sub-cellular systems is limited. A systems approach that integrates available experimental findings into a cohesive framework and connects sub-cellular systems influencing genotype to phenotype in HCM could address this knowledge gap and pave the way for identifying potential drug targets.

In many HCM variants, mutation-induced changes in myofilament calcium sensitivity and the resulting increase in sarcomere active force have been identified as the primary factor leading to hypertrophic remodeling of cardiomyocytes[17–19]. However, the involvement of multiple biochemical signal transduction modules, such as Ca^2+^, phosphoinositide 3-kinase/kinase B (PI3K/AKT), Ras/Raf/extracellular signal-regulated kinase 1/2 (ERK1/2), titin-associated proteins, calcineurin/nuclear factor of activated T-cells (CaN/NFAT), mammalian target of rapamycin (mTOR), growth factors, and Gq protein-coupled receptors in regulating the cardiomyocyte hypertrophic response to the altered force generation has been demonstrated by multiple studies [15, 17, 18]. In addition to the signaling modules, a few studies have underscored the remodeling of the cardiomyocyte metabolic network at the early stages of disease progression and its influence on cardiomyocyte hypertrophic growth and remodeling [14, 20–23]. According to these studies, in HCM, cardiomyocytes revert to a fetal metabolic profile, characterized by elevated glucose dependency and diminished oxidative capacity [14]. Deficiencies in ATP production, unbalanced reactive oxygen species (ROS) homeostasis, and dysfunctional nitric oxide synthases (NOS) activity are also re-ported in hypertrophic cardiomyopathy affecting cardiomyocyte growth and remodeling [20–23]. However, how cardiac signaling and metabolic networks, including their complex interplay, govern the genotype-to-phenotype relationship in hypertrophic cardiomyopathy remains unclear. In this study, we seek to use a systems biology modeling approach to close this gap and predict potential drug targets for HCM (see Fig. 1A).

**Figure 1:**
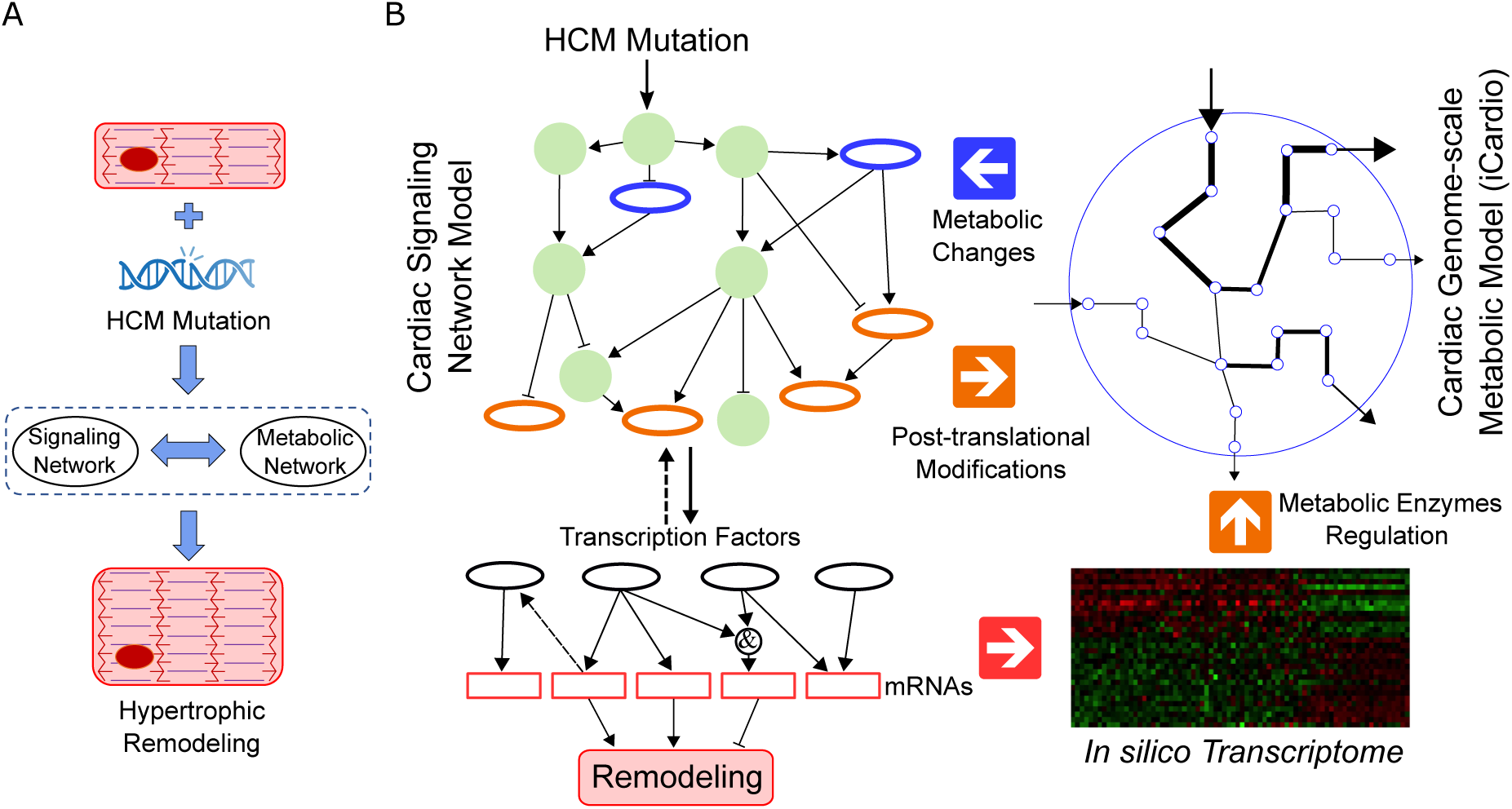
Systems modeling of genotype to phenotype in HCM. (A) Mapping HCM mutation in sarcomeric proteins to the hypertrophic remodeling of cardiomyocytes through signaling and metabolic networks. (B) The pipeline used to develop the systems biology model of HCM. This model connects the expanded version of the cardiomyocyte signaling network model to a genome-scale metabolic network model of the heart (iCardio [10] through a gene regulatory network model.

Our primary hypothesis is that in HCM, hypertrophic remodeling of cardiomyocytes is jointly regulated by both cardiac signaling and metabolic networks. If this hypothesis is valid, then molecular species that link these two cellular systems could act as potential drug targets to prevent or reverse hypertrophic cardiomyopathy. To test this hypothesis, we constructed a comprehensive model that connects sub-cellular systems in HCM, including cardiomyocyte signaling, metabolic, and gene regulatory networks. Previously, we had developed a signaling network model of cardiomyocyte morphological changes in familial cardiomyopathy based on experimental findings from preclinical research (e.g., murine data) [24]. There, we linked mutation-induced changes in sarcomere Ca^2+^ sensitivity to distinct morphological changes (i.e., cell area and elongation) of cardiomyocytes in hypertrophic and dilated familial cardiomyopathy [24]. In this study, we expanded our previously developed signaling network model to incorporate post-translational regulators (e.g., AMP-activated protein kinase: AMPK), transcriptional regulators (e.g., transcription factor EB: TFEB), and metabolites (e.g., ATP and ROS) that bridge cardiac signaling to the metabolic network. By coupling the expanded model to a heart-specific genome-scale metabolic network model (iCardio) [10], we reconstructed a computational systems biology model of HCM (see Fig. 1B) that incorporates crosstalk between cardiomyocyte signaling and metabolism. The iCardio model uses the TIDEs approach (Tasks Inferred from Differential Expression) to determine significant changes in cardiomyocyte metabolic functions associated with changes in mRNA expression. As opposed to the conventional gene-centered approach in standard gene enrichment like Gene Set Enrichment Analysis (GSEA), the TIDEs approach adopts a reaction-centered methodology [10]. The TIDEs approach accommodates both the stoichiometric balance of reactions and the complex interconnections among genes, proteins, and the reactions they catalyze to achieve metabolic functions with higher accuracy [10].

The developed HCM model links cardiomyocyte signaling, gene regulatory, and metabolic networks to simulate hypertrophic growth and remodeling of cardiomyocytes after the mutation-induced increase in sarcomere Ca^2+^ sensitivity in HCM. Utilizing this model, we aimed to address the following questions: How might ATP deficiency contribute to cardiomyocyte remodeling in HCM? Which regulatory reactions could control the transition from genotype to phenotype in HCM across different species or mutations? Which potential drug targets could prevent or reverse cardiomyocyte remodeling in HCM? The model predicted the potential impact of HCM-induced ATP deficiency on metabolic processes primarily associated with glutamine, as well as an elevation in cardiac apoptosis. Our findings also revealed shared (e.g., activation of PGC1*α* by AMPK, and FHL1 by titin) and context-specific mechanisms (e.g., regulation of Ca^2+^ sensitivity by titin in HCM patients) that control cardiomyocyte response across various contexts. We also predicted potential combination drug targets for HCM (e.g., mavacamten paired with ROS inhibitors) and evaluated their potential in preventing or reversing hypertrophic growth, apoptosis, and metabolic remodeling in cardiomyocytes.

## 2 RESULTS

According to previous experimental studies, both cardiomyocyte signaling and metabolic networks contribute in genotype to phenotype mechanisms in familial hypertrophic cardiomyopathy (HCM) [14, 15, 17, 20, 25]. In this study, we developed a computational systems biology model of hypertrophic cardiomyopathy (HCM) to mechanistically investigate the role of cardiomyocyte signaling and metabolic networks in governing the pathological remodeling processes of cardiomyocytes in HCM.

### 2.1 Predictive systems biology model of HCM

The complexity of signaling and metabolic pathways regulating cardiomyocyte remodeling complicates the identification of key mechanisms linking genotype to phenotype in HCM. To address this challenge, we developed a computational systems biology model of HCM that incorporates cardiomyocyte intracellular networks, including signaling, gene regulatory, and metabolic networks, as well as their crosstalk. Using a stochastic logic-based ODE approach [26], we simulated the impact of the mutation-induced increase in sarcomere Ca^2+^ sensitivity as the model input on the hypertrophic growth and remodeling of cardiomyocytes as the output. We used a stochastic differential equation approach to capture the extrinsic noise of molecular species, providing a more accurate model of cellular processes in cardiomyocytes [27]. In our model, we integrated three interaction types between cardiomyocyte signaling and metabolic networks including 1) transcriptional, and 2) post-translational regulation of metabolic enzymes, as well as 3) metabolic network feedback to the signaling through metabolites as shown in Fig. 1B.

To create the HCM model, we identified key metabolites, metabolic enzymes, and transcription factors linking cardiomyocyte signaling and metabolic networks by reviewing over 80 articles. We integrated these elements into our prior familial cardiomyopathy signaling model (36 species) [24], yielding an expanded HCM signaling network with 85 species and 187 reactions as displayed in Fig. 2A. Considering the steady-state outputs of the iCardio metabolic model, we integrated key AMPK-related pathways into the expanded signaling network to capture AMPK dynamics in our predictions. The expanded signaling network model incorporates a wide range of signaling components including environmental factors, receptors, upstream regulators, Ca^2+^ signaling regulators, signaling hubs, downstream regulators, metabolic mediators, and transcription factors forming a complex large-scale network. The 23 transcription factors, primarily governing metabolic enzyme genes and cardiomyocyte growth, bridge the signaling and gene regulatory networks as shown in Fig. 2A (cyan ovals). These transcription factors directly control the expression of 1078 genes, producing an *in silico* transcriptome. Feedback pathways were added from 49 genes to associated signaling nodes to model protein translation linked to these genes. We used the AND logic for modeling feedback to signaling nodes with multiple related genes, like PI3K/AKT. The *in silico* transcriptome was used as the input to the iCardio metabolic model that was built previously [10] using a genome-scale metabolic model of Homo sapiens (iHsa) [28] with tissue-specific protein data from the Human Protein Atlas (HPA) [29]. For metabolic enzymes regulated by both transcriptional and post-translational modifications, a linear function was used to combine the effects. Beyond the integrated interactions in the expanded signaling model, metabolic changes estimated by the iCardio model were also fed back into the signaling model.

**Figure 2:**
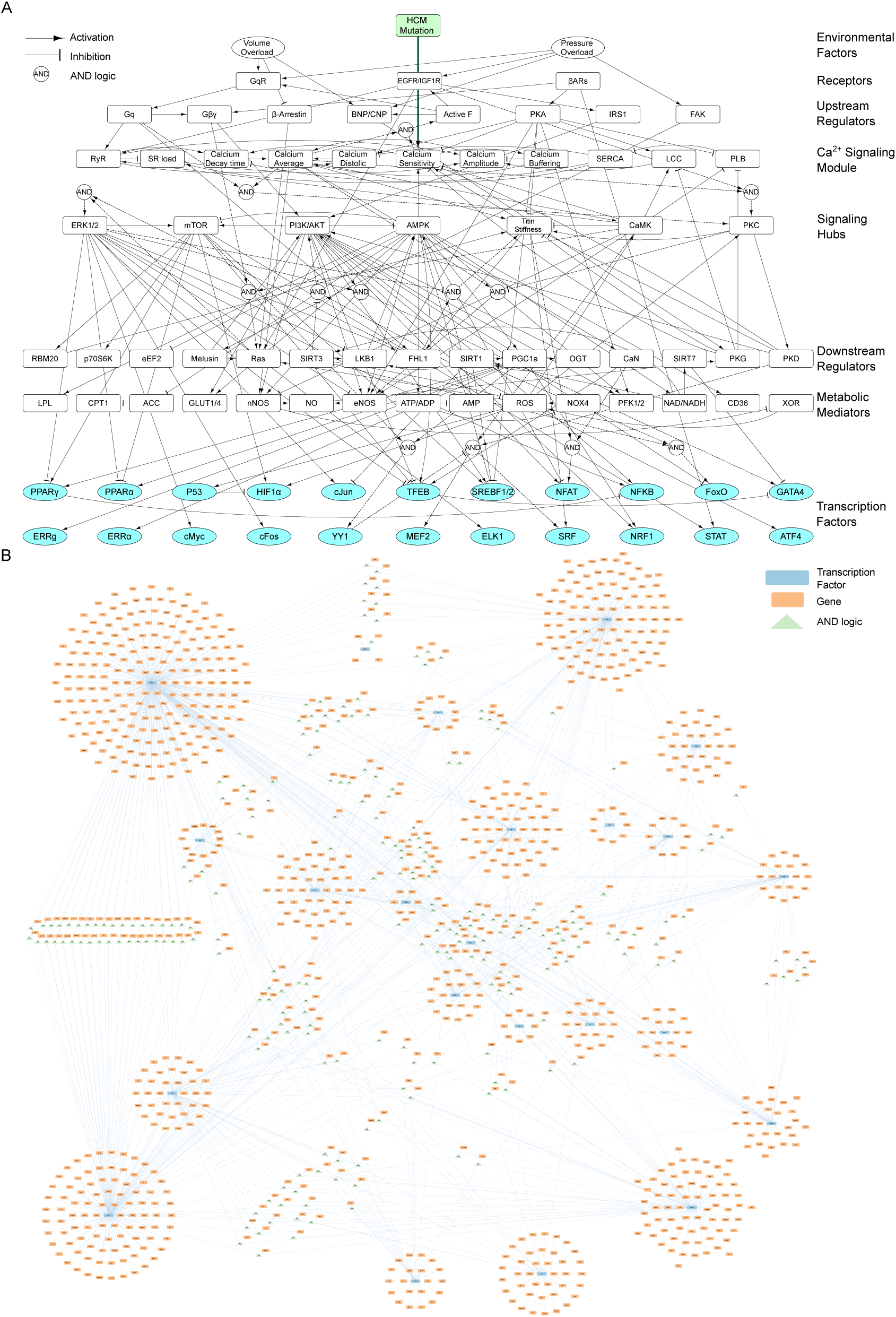
The systems biology model of HCM. (A) Schematic of expanded cardiomyocyte signaling network model in HCM. The model incorporates 85 signaling species connected through 187 reactions. (B) Gene regulatory network model of HCM linking 23 transcription factors (TFs) to their directly regulated 1078 genes. In a circular layout, genes regulated by one TF are positioned close to it, while those co-regulated by multiple TFs are associated with an AND gate (green triangle).

### 2.2 Model validation with HCM-specific data

To validate the model accuracy for HCM, we compared its predicted signaling activities and gene expressions at steady state to existing experimental data. Despite the prior validation of the original signaling model with HCM data [24], its extensive expansion with new species and multiple feedback required reassessment for accurate HCM predictions using independent studies.

We employed classified qualitative validation [11] to compare model predictions with experimental data. From 27 articles distinct from references used for model development, curated experimental data were sorted into three groups – decrease, no change, or increase – based on their statistical significance against study controls (see Supplementary Table 1). For model predictions, the HCM-induced activity of signaling nodes at steady-state were compared to their steady-state control with no HCM input. Nodes with non-significant changes (FDR > 0.05) were classified as no change, like nitric oxide (NO). As depicted in Fig. 3A, the model accurately predicted 83% of the experimental outcomes linked to 32 signaling nodes in the HCM context. For nodes like SERCA with varying observations, all these observations were equally weighted in the validation analysis (e.g., each SERCA observation was given half-weight). Additionally, since HCM-induced changes in the metabolic network model (e.g., change in ATP synthase and levels) typically span a longer time frame than alterations in signaling species, we assessed model predictions in two stages: before and after integrating feedback from the metabolic to the signaling network. Hence, for certain elements like AMPK and acetyl-coenzyme A carboxylase (ACC), the model predicted divergent directions of changes across these states. Such different predictions are depicted in Fig. 3A and incorporated into the validation analysis. For certain elements like AMPK and acetyl-coenzyme A carboxylase (ACC), the model predicted divergent directions of changes across these states which factored into the validation analysis.

**Figure 3:**
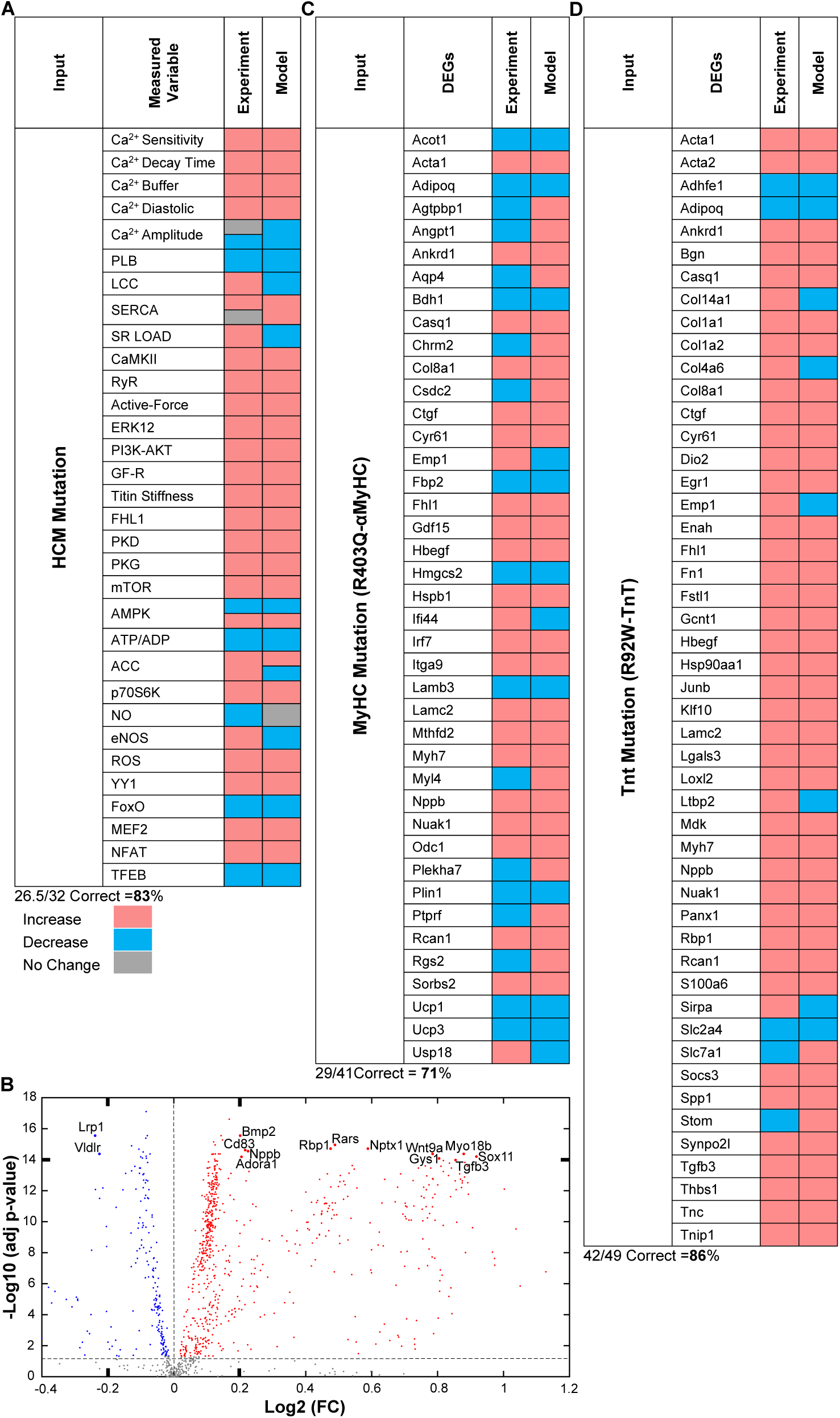
The model accurately predicts classified qualitative changes in the activity of signaling nodes and cardiomyocyte transcriptome measured by independent experimental studies. (A) Comparison of model predictions with experimental data of 32 signaling nodes from literature. (B) The *in silico* transcriptome indicates upregulated and downregulated differentially expressed genes in HCM. (C) Comparison of model-predicted DEGs with 41 DEGs in αMyHC-mutant mouse hearts (R403Q-αMyHC)[30]. (D) Comparison of model-predicted DEGs with 49 DEGs in TnT-mutant mouse hearts (R92W-TnT)[30].

Employing the model, we then generated the *in silico* transcriptome of HCM. The *in silico* transcriptome (Fig. 3B) indicates both upregulated and downregulated differentially expressed genes (DEGs) within the HCM context (see Supplementary Table 1). As the model uses default parameters, the estimated mRNA fold changes might not be sufficiently precise for determining significance levels, as practiced in some studies [31]. Thus, DEGs were determined using only an FDR threshold of < 0.05. To validate the *in silico* transcriptome predicted by the model, We employed two published mouse models of HCM [30]. Using classified qualitative validation [11], DEGs from young mice with R403Q-αMyHC and R92W-TnT mutations were compared to the *in silico* transcriptome, as shown in Figures 3C and D. We chose these HCM mouse models because they showcase the early HCM stages where remodeling in cardiomyocyte intracellular networks is more readily distinguishable [30]. The R403Q-MyHC mutation in the α-myosin motor domain of mice, which some studies suggest displays enhanced ATPase activity as a molecular gain-of-function mutation, results in left ventricular hypertrophy and heart failure [30]. The R92W-TnT mutation in cardiac troponin T leads to increased Ca^2+^ sensitivity and is associated with cardiac fibrosis, sudden cardiac death early in life, and lower levels of hypertrophy compared to R403Q-αMyHC mutation [30]. Out of 223 mRNAs that were differentially expressed in R403Q-αMyHC mutation, 41 mRNAs were common with model-predicted DEGs where the model correctly predicted changes in 71% of them (see Fig. 3C). For R92W-TnT mutation, 265 mRNAs were differentially expressed compared with their controls [30] and the model correctly predicted 86% of 49 common DEGs.

In summary, the HCM model was validated against existing experimental data in the HCM context, achieving a high prediction accuracy (83%) for data linked to signaling nodes as well as predicting qualitative changes in differentially expressed genes of R403Q-αMyHC (71%) and R92W-TnT (86%) mutant mouse models.

### 2.3 HCM-driven ATP deficiency could control cardiomyocyte metabolic shift and apoptosis

HCM mutations could change calcium sensitivity in sarcomeres, influencing not only cardiomyocyte contraction but also multiple cellular activities that promote cardiac cell growth and remodeling. Thus, using the HCM model, we predicted potential changes in the metabolic functions of cardiomyocytes associated with HCM. As illustrated in Figures 4A and B, the model anticipated a decrease in palmitate intake and fatty acid metabolism, while expecting an increase in the creatine kinase system and carbohydrate metabolism. Taking into account all significant metabolic changes predicted by the model (refer to supplementary Table 2), HCM mutation could lead to reduced ATP synthase accompanied by a metabolic shift from mainly relying on fatty acids for energy to using carbohydrates and other sources. These predictions align with past experimental research conducted on HCM mouse models and HCM patients [30, 32–35].

**Figure 4:**
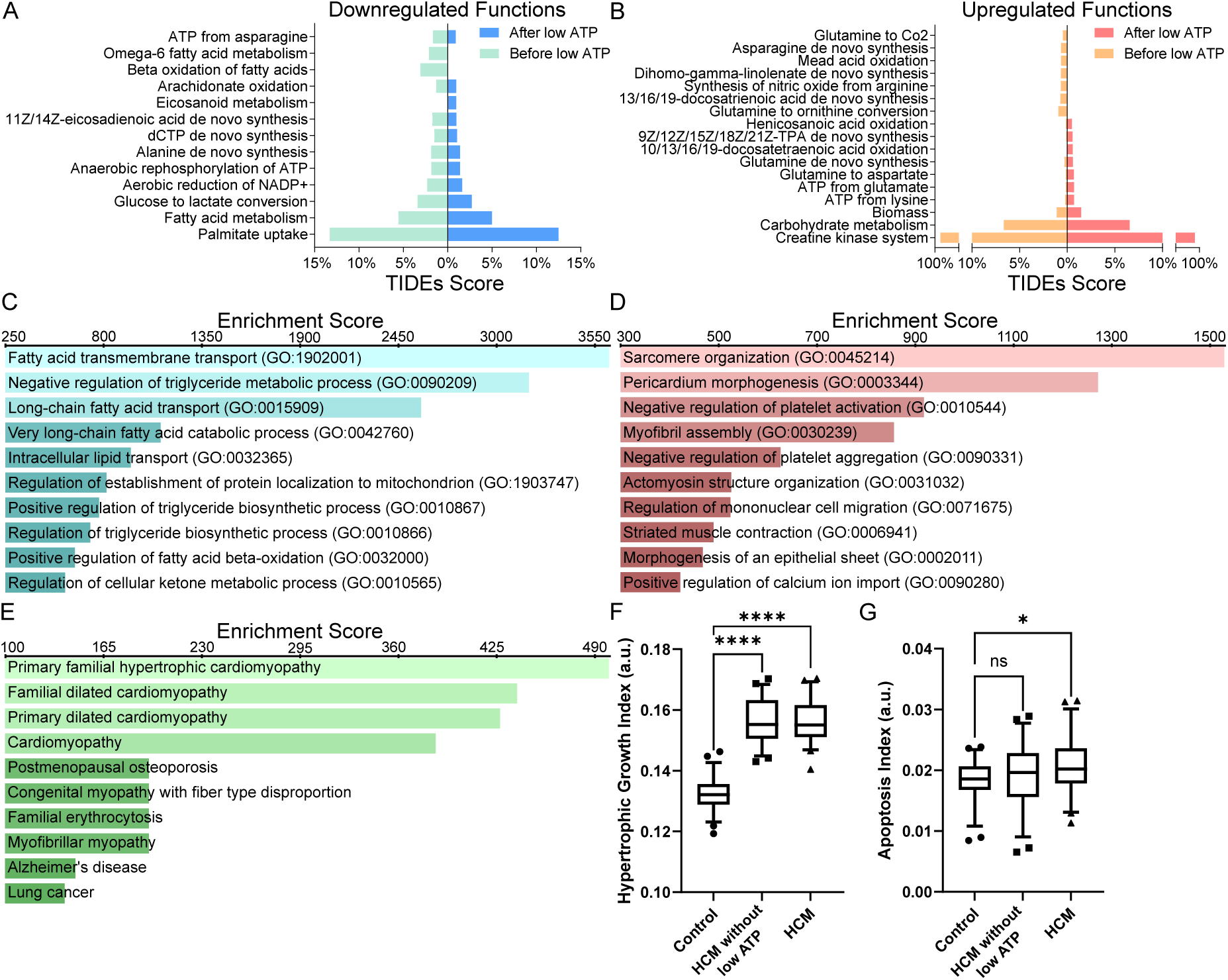
The model predicts significant remodeling in metabolic functions and sarcomere structure/organization in HCM. (A) Top significant (p-value<0.025) downregulated and (B) upregulated metabolic functions in HCM predicted by the model using the TIDEs approach before and after applying feedback from the metabolic network model to the signaling (i.e., lower ATP synthase and level). The TIDEs approach identifies metabolic functions linked to DEGs by applying gene expression fold changes using gene-protein-reaction (GPR) rules to weight network reactions. TIDEs’ task scores are determined by averaging the reaction weights within a task [10]. (C) Gene Ontology analysis results for downregulated and (D) upregulated DEGs predicted by the model. (E) Results of ClinVar enrichment analysis for upregulated DEGs predicted by the model. (F) HCM-induced changes in hypertrophic growth and (G) apoptosis indexes predicted by the model. The Whiskers show 5 and 95 percentiles. Fifty *in silico* samples for each condition were used for statistical analysis (ordinary one-way ANOVA followed by Dunnett’s multiple comparison test with single pooled variance: * p-value<0.05;**** p-value<0.0001). All analyses in (C-E) were conducted by Enrichr, a web-based enrichment analysis platform, and combined scores computed by multiplying the log of the p-value from the Fisher exact test by the z-score of the deviation from the expected rank were used for ranking biological functions and diseases. [36].

Although the expanded signaling network model captured the HCM-induced imbalance between ATP supply and demand [37] by predicting a decrease in ATP level, we further adjusted the Ymax of ATP level to 80% of its initial level [38] to represent the advanced stages of the disease. After the adjustment, we ran the model to a new steady state for further analyses in subsequent sections. To assess the impact of deficiencies in ATP synthase on cardiomyocyte metabolism in the HCM context, we illustrated the predicted changes in metabolic functions for both states. As shown in Fig. 4A, deficiencies in ATP production diminish the degree of downregulation in metabolic functions to partially compensate for the low ATP level. On the other hand, as shown in Fig. 4B, the degree of upregulation for metabolic functions associated with glutamine significantly alters after lowering the ATP level. To evaluate how ATP synthase deficiencies affect cardiomyocyte metabolism in HCM, we presented the predicted metabolic changes before and after adjusting ATP levels. As illustrated in Fig. 4A, ATP production deficiencies could partially offset the metabolic downregulation to counterbalance the reduced ATP. The model also predicted significant upregulation in glutamine-related metabolism after ATP level reduction (Fig. 4B). For example, following the reduction in ATP production, the model predicted a substantial upregulation in the conversion of glutamine to aspartate, de novo synthesis of glutamine, and ATP production from glutamate. Conversely, the previously significant HCM-induced enhancement of the conversion pathways from glutamine to carbon dioxide (CO2) and glutamine to ornithine became insignificant after applying ATP feedback. These alterations might suggest an intensification of the glutaminolysis process within cardiomyocytes, which potentially exerts cardioprotective impacts by compensating for the diminished ATP production [39]. The model predicted two other noteworthy changes following ATP level reduction: an increase in biomass production and the nullification of the HCM-induced increase in nitric oxide synthesis from arginine (Fig. 4B).

Moreover, using the *in silico* transcriptome and Gene Ontology (GO) analysis [40] via Enrichr [36], a web-based enrichment analysis platform, we determined key biological processes affected by the HCM mutation. The GO analysis revealed downregulation in processes like fatty acid transport, and negative regulation of the triglyceride metabolic process (Fig. 4C). In contrast, processes like sarcomere organization, myofibril assembly, muscle contraction, and positive regulation of Ca^2+^ import were upregulated in the HCM setting (Fig. 4D). Using Enrichr for ClinVar enrichment analysis (V. 2019) [41], we also identified related human diseases associated with upregulated mRNAs in *in silico* transcriptome. As anticipated, familial hypertrophic cardiomyopathy scored highest, followed by other forms of cardiomyopathy. Furthermore, given the importance of cardiomyocyte hypertrophy and apoptosis in heart failure [42], we developed two indexes representing mRNAs associated with the regulation of cardiomyocyte hypertrophic growth and apoptosis, the model predicted a marked rise in cardiomyocyte hypertrophic growth in both HCM states (Fig. 4F). The model predicted no significant differences in cardiomyocyte apoptosis between the control and HCM without considering the feedback from reduced ATP levels. However, simulating the ATP synthase deficiency caused by HCM resulted in a slight but notable increase in the apoptosis index, suggesting ATP deficiency’s role in adverse cardiomyocyte remodeling in HCM (Fig. 4G).

To summarize, the HCM model predicted significant alterations in cardiomyocyte metabolic functions, emphasizing ATP synthase deficiency and a transition from fatty acids to carbohydrate metabolism. GO analysis of *in silico* transcriptome predicted increased sarcomere organization and contraction as well as decreased fatty acid transport in HCM. Additionally, the model indicated major shifts in glutamine-related metabolism and an uptick in cardiac apoptosis due to ATP synthase deficiency.

### 2.4 HCM model reveals shared and context-specific regulatory reactions controlling genotype to phenotype in various HCM contexts

Identifying key regulatory reactions controlling cardiomyocyte response in HCM can deepen our understanding of genotype-phenotype mechanisms, paving the way for better therapeutic strategies. Using the HCM model, we performed a global sensitivity analysis (via the Morris method [43]) by adjusting the weight parameter (W) to identify the main regulatory reaction in the HCM expanded signaling network controlling cardiomyocyte hypertrophic growth and apoptosis. In Figures 5A and B, each dot indicates the Morris index (μ*) and standard deviation (σ) for each reaction. Orange dots represent reactions with monotonic effects on the chosen model output (hypertrophic growth and apoptosis), while blue stars denote those with non-monotonic impacts. Given the absence of a standard criterion for the Morris index (μ*) [43], we set a threshold of 0.01, leading to the identification of 25 key reactions influencing hypertrophic growth and six major reactions affecting cardiomyocyte apoptosis in HCM. According to sensitivity analysis results, reactions that control the activity of transcription factors like STAT by mTOR, SRF by titin, TP53 by ROS, and GATA4 by SIRT7 linearly influence cardiomyocyte hypertrophic growth. Moreover, the significant standard deviation (σ) of the interaction between ATP and ROS mediated by NOX4 suggests a complex, non-linear regulation of hypertrophic growth by ATP-related ROS production, closely tied to other HCM network reactions. In terms of cardiomyocyte apoptosis, key regulatory reactions include TP53 activation by ROS, ROS modulation by NOX4 and SIRT3, STAT activation by mTOR and SRF by titin, and FoxO inhibition by PI3K/AKT.

**Figure 5:**
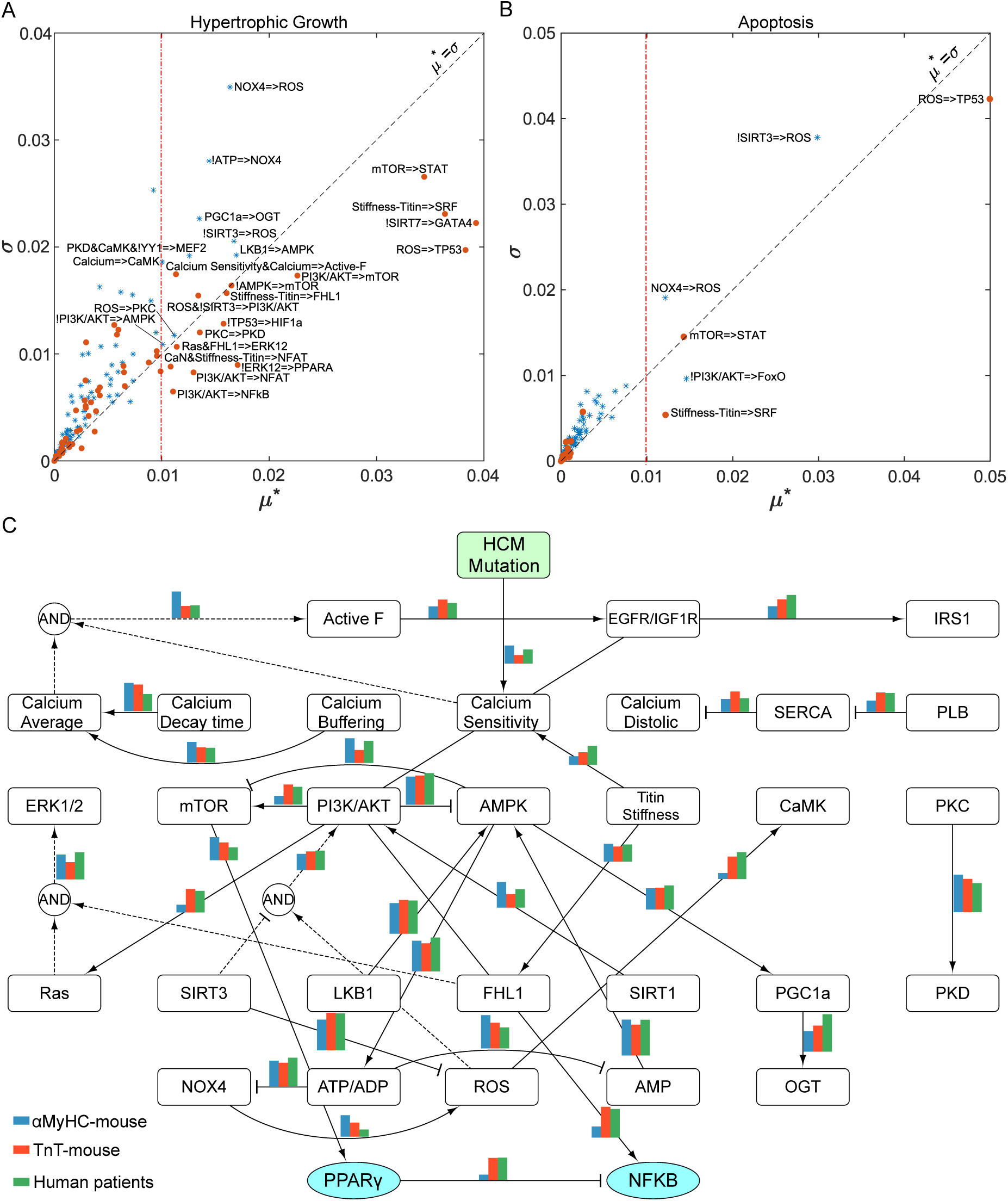
Morris global sensitivity analysis identifies major reactions regulating cardiomyocyte phenotype in different contexts. Important and less-important reactions in the HCM network regulating (A) the hypertrophic growth and (B) apoptosis of cardiomyocytes were identified based on the Morris index (μ*) larger or smaller than 0.01 (arbitrary criteria set by the designer), respectively. (C) Context-dependent regulatory reactions governing the cardiomyocyte response in three HCM contexts of mouse R403Q-αMyHC, mouse R92W-TnT, and human HCM patients. The top 20 reactions in each context are illustrated with three bar indicators comparing the normalized Morris indexes of each reaction.

Considering the significant variability in cardiomyocyte phenotypic responses to HCM mutations and differences between mouse and human HCM models, identifying major mechanisms controlling cardiomyocyte responses in specific HCM contexts is crucial but difficult through experimental research. To tackle this challenge, we used transcriptomic data from non-obstructive R403Q-αMyHC and R92W-TnT HCM mutations in mouse models, as well as human transcriptomic datasets (GSE36961: mRNA, GSE36946: miRNA) [30, 32], to identify the key context-specific reactions that modulate cardiomyocyte response in HCM. Using the accuracy of the model in predicting context-specific transcriptomes as the output of the Morris sensitivity analysis, we identified the top 20 key reactions for each context. Fig. 5C displays three bar indicators comparing the normalized Morris indexes across three studied contexts. Results indicated that cardiomyocyte response in HCM is directed by a mix of shared and context-specific reactions. Shared reactions across the three contexts include AMPK activating PGC1*α*, titin activating FHL1, and AMPK regulating ATP/ADP levels, with AMPK itself being activated by LKB1 and inhibited by PI3K/AKT.

Based on model predictions, interactions such as ROS production by NOX4, PKD activation by PKC, the activation of PPAR*γ* by mTOR, and the regulatory reactions linking Ca^2+^ transients to the sarcomere active force are central in controlling the cardiomyocyte response for the αMyHC mutation. On the other hand, for the TnT mutation, the model identified interactions like regulation of Ca^2+^ diastolic level by PLB through SERCA, NFKB regulation by PPAR*γ* and PI3K/AKT, CaMK activation by ROS, and activation of Ras through growth factor receptors as major regulatory reactions. In HCM patients, the model suggested a combination of major reactions in αMyhC and TnT contexts. However, the model predicted that some reactions, including regulation of Ca^2+^ sensitivity by titin, IRS1 activation by growth factor receptors, and OGT regulation by PGC1*α*, are more pronounced in HCM patients.

In brief, our model predicted that the activity of transcription factors such as STAT, SRF, GATA4, TP53, and FoxO, are key regulators of cardiomyocyte hypertrophy and apoptosis in HCM, aligning with prior experimental studies [44–48]. Furthermore, the model revealed the presence of shared and context-specific mechanisms that control genotype to phenotype transition in HCM, potentially accounting for the heterogeneity in the cardiomyocyte response observed in different HCM mutations or experimental models.

### 2.5 HCM model predicts potential combination therapies preventing/reversing HCM phenotype

Recently, mavacamten, a cardiac myosin inhibitor that decreases actin-myosin ATPase activity and sarcomere hypercontractility in HCM, exhibited noteworthy effects in treating patients diagnosed with obstructive hypertrophic cardiomyopathy [49]. However, the complexity of mechanisms linking genotype to phenotype in familial hypertrophic cardiomyopathy hinders the prediction of new drug targets for this cardiac muscle disorder [50]. Considering the increasing trend in clinical trials for combination therapies [51]—attributed to their enhanced potential in addressing intricate cardiovascular conditions like heart failure [52]—we hypothesized that these therapies might also offer a viable treatment option for HCM. Accordingly, we performed a combinatory perturbation analysis to identify potential drug targets for HCM. In this analysis, we centered our focus on combined inhibitions of model species, drawing from our previous research [24] that showcased the higher effectiveness of drug targets in HCM when inhibiting signaling nodes together.

Upon examining node pair inhibitions by modifying the Ymax parameter from 1.0 to 0.1, we assessed their effects on hypertrophic growth and apoptosis indexes. We determined the effectiveness of drug combinations against HCM by comparing fold change ratios of indexes in the “HCM with drug” to the “control with drug” conditions. As depicted in Fig. 6A, several combined inhibitory targets, indicated by red dots, notably decreased both indexes, suggesting their potential to prevent or reverse the HCM phenotype. The model results suggested that reducing Ca^2+^, Ca^2+^ sensitivity, and active force, in conjunction with inhibition of targets such as ATP, ROS/NOX4, TP53, and AMPK, could effectively counteract the HCM-induced changes in cardiomyocyte apoptosis and hypertrophic growth. To assess the significance of these pharmacological interventions on cardiomyocyte hypertrophic growth and apoptosis, we selected some drug targets paired with Ca^2+^ sensitivity and showing high potential. While the combination of AMPK hyperactivation and Ca^2+^ sensitivity reduction did not rank among the top drug interventions, we also included it in our analysis to examine the potential influence of metformin, a widely used AMPK activator for diabetes treatment, on potential drugs targeting Ca^2+^ sensitivity in hypertrophic cardiomyopathy. Numerous HCM mutations increase the sarcomeres’ Ca^2+^ sensitivity. As expected, the model identified Ca^2+^ sensitivity reduction as a viable intervention. However, we were more interested in drug targets enhancing the efficacy of this intervention. Thus, for further analysis, we focused on drug targets paired with Ca^2+^ sensitivity demonstrating higher efficacy. Given the strong relationship between Ca^2+^ sensitivity and active force, the results of this analysis could also apply to drugs that decrease sarcomere active force like mavacamten.

**Figure 6:**
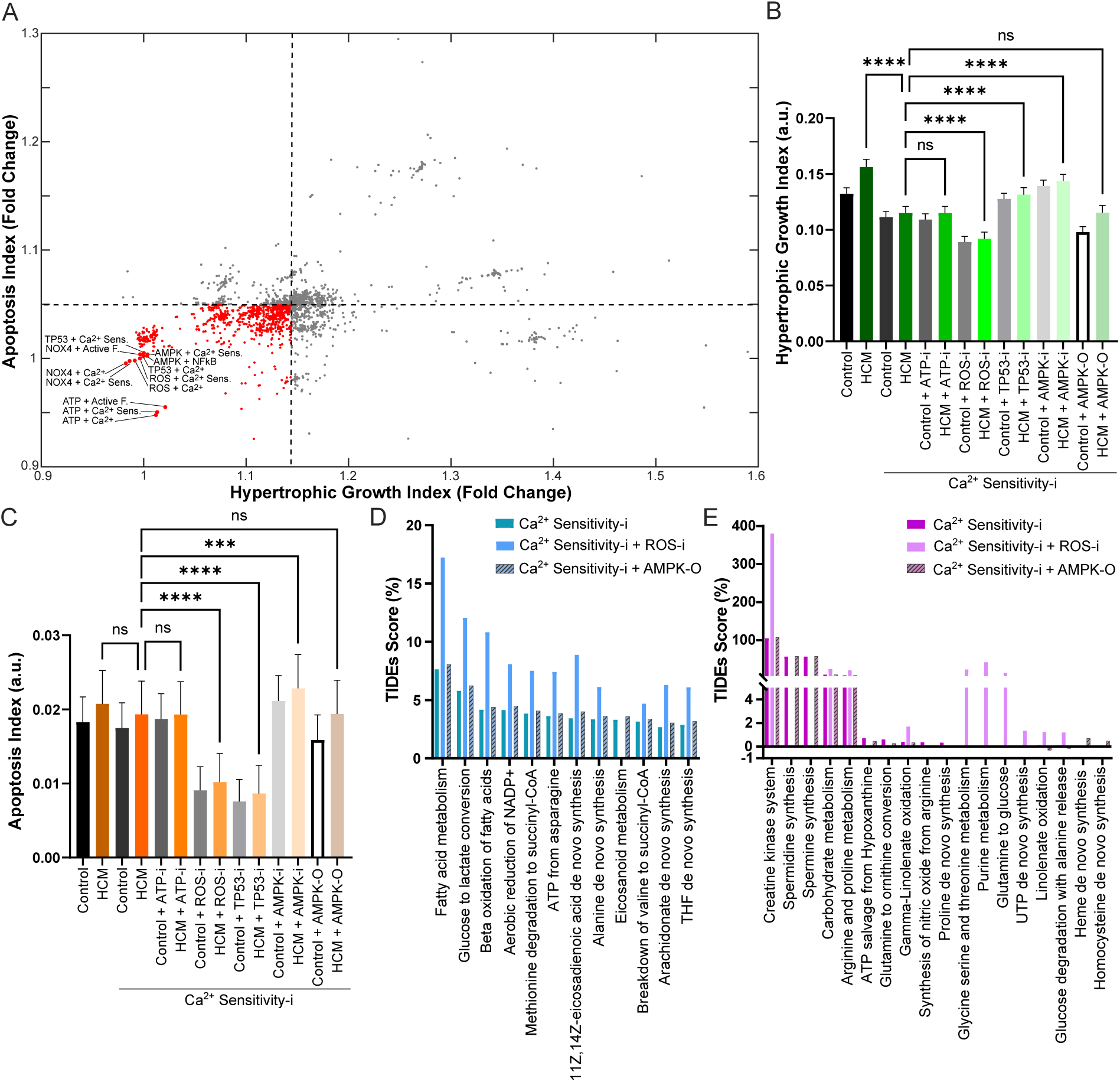
The model predicts potential combination drug targets in the HCM context. (A) Efficacy of combination pharmacotherapy on hypertrophic cardiomyopathy. Data represent all pairwise combinations of inhibiting (Ymax = 1 to 0.1) node activity and their impact on hypertrophic growth and apoptosis indexes in HCM. Red points indicate potential drug targets preventing/reversing cardiomyocyte remodeling. (B) Model-predicted hypertrophic growth and (C) apoptosis indexes (Mean*±*SD) in selected drug targets in the HCM context. Fifty *in silico* samples for each condition were used for statistical analysis (ordinary one-way ANOVA followed by Tukey’s multiple comparison test with single pooled variance:*** p-value<0.001;**** p-value<0.0001). (D) Model-predicted metabolic functions that were upregulated and (E) downregulated in analyzed drug interventions.

After analyzing hypertrophic growth and apoptosis indexes following perturbation of selected drug targets (inhibition: Ymax=1.0 to 0.5; hyperactivation: Ymax=1.0 to 1.5), we detected a significant reduction in the hypertrophic growth index (Fig. 6B) and no significant variation in the apoptosis index (Fig. 6C) when *in silico* HCM cardiomyocytes were simulated with a Ca^2+^ sensitivity reduction, compared to untreated HCM cardiomyocytes. When paired with Ca^2+^ sensitivity reduction, a decrease in ATP level did not impact hypertrophic growth or apoptosis compared to the reference condition of HCM cardiomyocytes with Ca^2+^ sensitivity reduction. Conversely, combining ROS inhibition with Ca^2+^ sensitivity reduction notably reduced both indexes. Adding a TP53 inhibitor decreased the apoptosis index but raised the hypertrophic growth index. AMPK inhibition raised both indexes, while AMPK hyperactivation showed no significant effect, suggesting a nonsignificant impact of metformin on the efficacy of Ca^2+^ sensitivity reduction at the cardiomyocyte level.

Using the model, we also evaluated the effects of certain drug targets on cardiomyocyte metabolism. In addition to the Ca^2+^ sensitivity reduction, we explored its pairing with ROS inhibition due to its potential in reducing hypertrophic growth and apoptosis indexes, and the hyperactivation of AMPK given its crucial role in cardiomyocyte metabolic regulation. As depicted in Fig. 6D, all these interventions promoted reverse metabolic remodeling in cardiomyocytes compared to the HCM condition by upregulation of fatty acid metabolism, glucose-lactate conversion, beta-oxidation of fatty acids, aerobic reduction of NADP+, and ATP generation from asparagine. Among the evaluated drug interventions, the combination of Ca^2+^ sensitivity reduction and ROS inhibition notably enhanced the metabolic functions. On the other hand, combining AMPK hyperactivation with Ca^2+^ sensitivity reduction didn’t substantially augment the outcomes of Ca^2+^ sensitivity reduction alone. Regarding the downregulated metabolic functions after studied interventions (Fig. 6E), the decrease in creatine kinase system, carbohydrate metabolism, and arginine and proline metabolism recorded the highest TIDEs scores across all treatments. While certain reductions, like in the creatine kinase system and carbohydrate metabolism, effectively countered HCM-induced changes, functions linked to glutamine (elevated in the HCM condition, as per Figure 4B) remained relatively unchanged after interventions, suggesting only a partial metabolic restoration. Model findings also highlighted metabolic changes exclusive to the paired inhibition of Ca^2+^ sensitivity and ROS, such as reductions in glycine serine and threonine metabolism, purine metabolism, and the conversion of glutamine to glucose. Refer to supplementary Table 2 for all significant metabolic function changes related to the above conditions.

In summary, using the HCM model, we identified potential combined drug targets for HCM and evaluated their effects on cardiomyocyte hypertrophic growth, apoptosis, and metabolic remodeling. The model predicted promising therapeutic targets with the joint inhibition of Ca^2+^ sensitivity/active force and ROS. It also indicated that metformin wouldn’t diminish the effectiveness of drugs reducing Ca^2+^ sensitivity/active force like mavacamten at the cardiomyocyte level.

## 3. DISCUSSION

Here, we have developed a cardiomyocyte-level systems biology model for familial hypertrophic cardiomyopathy (HCM) to study the effects of HCM mutations on sarcomeric proteins on cardiomyocyte phenotype. The model integrates three cellular networks including signaling, metabolic, and gene regulatory networks, and predicts HCM-driven cardiomyocyte growth and remodeling. We validated the HCM model against experimental data, curated from previous studies, indicating changes in the activity of signaling species in the HCM context, transcriptomes of two HCM mouse models (R403Q-αMyHC and R92W-TnT), and altered metabolic functions in HCM. The model suggested a shift in major metabolic substrates and revealed how HCM-induced ATP synthase deficiency impacts metabolic functions. By defining two indexes representing hypertrophic growth and apoptosis of cardiomyocytes, the model predicts cardiomyocyte phenotypic changes in HCM. The Morris global sensitivity analysis spotlighted key regulatory reactions and pathways controlling genotype to phenotype in different HCM contexts including two mouse HCM mutations in αMyHC (e.g., ROS production by NOX4, and activation of PPARγ by mTOR) and TnT proteins (e.g., NFKB regulation by PPARγ and PI3K/AKT, and CaMK activation by ROS) as well as HCM patients (e.g., regulation of Ca^2+^ sensitivity by titin and IRS1 activation by growth factor receptors). Finally, using a combined perturbation analysis, the model predicted some potential combination drug targets (e.g., joint inhibition of Ca^2+^ sensitivity and ROS) to prevent or reverse the HCM phenotype.

### 3.1 Metabolic remodeling in HCM

Metabolic remodeling is central to cardiomyocyte changes in hypertrophic cardiomyopathy [53]. Under physiological conditions, the heart predominantly relies on fatty acid oxidation for energy production. However, in HCM, an increase in the energy demand and mutation-induced changes in cardiomyocyte signaling and gene regulation lead to a metabolic shift towards increased glucose utilization and reduced fatty acid oxidation [54]. This transition causes inefficient ATP generation, inducing an energy shortage that can undermine myocardial contractility and cardiac performance [55]. Accumulation of metabolites like lactate and ROS in HCM can further intensify pathological heart remodeling [56]. These metabolic changes can intertwine with pivotal signaling routes governing myocardial growth. For instance, dysregulation of AMPK in HCM could affect cardiomyocyte hypertrophic growth [57].

Using the HCM model, we predicted metabolic function shifts in line with experimental findings [34, 54]. The model predicted a decline in palmitate absorption and fatty acid metabolism while projecting increases in the creatine kinase system and carbohydrate metabolism as primary metabolic alterations in HCM cardiomyocytes. Considering the observed association between cardiac palmitate uptake and the expression of fatty acid translocase (FAT/CD36) and fatty acid binding protein (FABPpm) [58], the predicted decline in palmitate uptake suggests a corresponding reduction in fatty acid metabolism. The model also predicted a slight reduction in glucose-to-pyruvate conversion and a larger decrease in glucose-to-lactate conversion, indicating reduced pyruvate-to-lactate conversion in early HCM stages. This response may be linked to the model-predicted decline in NADH/NAD+ ratio, given that prior research has highlighted the relationship of NADH/NAD+ ratio with the pyruvate to lactate flux [59].

The Creatine Kinase (CK) system, a primary energy buffer for the heart, responds rapidly to increased ATP needs, especially when the heart’s demand increases [60]. The model prediction of a substantial upregulation in the CK system might reflect an early cardiomyocyte adaptation to heightened ATP demand from an HCM mutation-driven increase in sarcomere active force. However, studies on heart failure [60, 61] and certain HCM patients (e.g., Arg403Gln mutation) [62] reported reduced CK activity, suggesting an ATP synthase deficiency. This emphasizes the role of CK in HCM mutation-induced metabolic remodeling of cardiomyocytes and invites more exploration into its therapeutic potential. The model also predicted a significant increase in carbohydrate metabolism which is in line with the experimental observations showing the “fetal metabolic profile” with an increased reliance on carbohydrate sources in HCM [63, 64].

### 3.2 ROS and hypertrophic cardiomyopathy

In HCM pathogenesis, Reactive Oxygen Species (ROS) play a pivotal role [65]. ROS, generated from cellular metabolism, can cause cell damage and inflammation when their concentrations surpass the inherent antioxidant defenses of cells, resulting in oxidative stress [66]. In HCM, elevated ROS levels have been linked to myocardial hypertrophy, fibrosis, and cardiac dysfunction, suggesting the therapeutic potential of ROS inhibitors for HCM [65]. ROS inhibitors can reduce oxidative stress by either scavenging the produced ROS or inhibiting their production. Preclinical studies have shown that the use of ROS inhibitors can attenuate myocardial hypertrophy and fibrosis, improve cardiac function, and potentially slow the progression of HCM [67]. Certain ROS inhibitors have shown anti-inflammatory and anti-apoptotic effects [68, 69], potentially mitigating the HCM-related pathological remodeling. While ROS inhibitors show promise for HCM treatment, challenges persist. For example, the timing and duration of ROS inhibition are crucial, as long-term ROS inhibition might interfere with the physiological roles of ROS, potentially causing side effects [70]. Moreover, given the heterogeneous nature of HCM and complex ROS signaling, ROS inhibition might affect individuals differently [22, 65, 67], potentially contributing to the limited effectiveness of general antioxidant treatments for patients with heart disease [71].

In this study, we identified ROS and its upstream and downstream reactions as major regulators of cardiomyocyte hypertrophic and apoptotic response in HCM. Sensitivity analyses (Figs. 5A and B) highlighted that while ROS-driven TP53 activation significantly affects cardiomyocyte hypertrophy and apoptosis monotonically, reactions regulating ROS generation and scavenging exhibited nonlinear, variable impacts on the cardiomyocyte phenotype. Prior research suggests that ROS-mediated TP53 activation can drive cardiomyocyte metabolic remodeling [72] and transition from adaptive cardiac hypertrophy to maladaptive cardiac dysfunction via pro-apoptotic and anti-angiogenesis functions of TP53 [47]. Our context-specific analysis highlighted considerable variability in reactions at both upstream (e.g., NOX4 => ROS) and downstream of ROS (e.g., ROS => CaMK) across different HCM mutations and species (refer to Fig. 5C). Past research has shown mixed effects of NOX4 in cardiac disease models [73]. Some studies suggest NOX4 has a protective role against cardiac hypertrophy and stress-induced cardiac remodeling [74], while others link it to negative outcomes due to increased ROS production and cell damage [75]. ROS-dependent oxidation of CaMKII is also reported in various cardiac disease models contributing to calcium handling abnormalities and impaired contraction [76], myocardial injury and inflammation [71], and arrhythmia [77]. We also predicted that jointly inhibiting ROS and Ca^2+^ sensitivity could prevent or reverse the HCM phenotype in cardiomyocytes. While our findings support targeting ROS as a potential therapeutic strategy for HCM, the complex and nonlinear impact of ROS on cardiomyocytes underscores the importance of using context-specific models in developing ROS-targeted therapies.

### 3.3 Heterogeneity in HCM

Familial hypertrophic cardiomyopathy is a heterogeneous disease, both genetically and clinically [50, 78, 79]. Genetically, HCM is primarily linked to mutations in genes encoding sarcomeric proteins, but non-sarcomeric gene mutations are also found [50]. Clinically, the HCM phenotype is highly variable, from patients with no symptoms to those experiencing severe heart complications like diastolic dysfunction, obstructive HCM, and heart failure [80]. The degree of hypertrophy, pattern of ventricular remodeling, presence of myocardial fibrosis, and occurrence of arrhythmias can vary significantly among HCM patients [79]. Disease progression is also unpredictable, with some individuals remaining stable for years while others rapidly deteriorate by heart failure or sudden cardiac death [81]. These clinical variations and diverse mutations in HCM complicate drug development, as a treatment effective for one patient might not work for another [82]. Therefore, understanding the molecular and cellular mechanisms underlying HCM heterogeneity can promote the development of effective strategies for treating HCM patients.

We used the HCM model to mechanistically investigate mechanisms that may underpin HCM heterogeneity across different mutations. Our results, shown in Fig. 5, revealed some mutation-specific molecular reactions that may enlighten some discrepancies observed in experimental models with different HCM mutations [30]. For example, Vakrou and colleagues [30] suggested a higher upregulation of endothelin-1 signaling in mice with TnT mutation, whereas they predicted a more pronounced downregulation of calcium signaling in mice with MyHC mutation. Endothelin-1 signaling includes activation of Gq/G*βγ*/Ca^2+^/PKC [83], and EGFR/Ras pathways [84]. While the HCM model predicted a higher score for both upstream and downstream reactions of EGFR/Ras signaling in TnT mutants, no significant difference was observed for Gq/G*βγ*/Ca^2+^/PKC pathway between TnT and MyHC mutants. This finding suggests that EGFR/Ras signaling may contribute to the observed differences between TnT and MyHC mutants in the context of hypertrophic cardiomyopathy. In terms of calcium signaling, the model predicted lower scores for reactions such as PLB-SERCA, SERCA-Ca^2+^ diastolic, and ROS-CaMKII in MyHC mutants suggesting a reduced influence of calcium signaling in regulating the early HCM phenotype in MyHC mutants compared to TnT mutants. The difference in NFkβ regulation between MyHC and TnT mutants was also observed in HCM model results. While short-term NFkβ activation can protect the heart during acute low oxygen conditions and reperfusion, prolonged activation might lead to heart failure by causing chronic inflammation [85]. Accumulating evidence supports the critical role of NFkβ and chronic inflammation in hypertrophic cardiomyopathy [86]. Kuusisto et al. [87] found varying degrees of inflammatory cell presence and NFkβ activation in HCM patients’ hearts, with a strong correlation between inflammation and cardiac fibrosis. High NFkβ baseline levels predicted progressive heart failure in mildly symptomatic or asymptomatic HCM patients over a 10-year follow-up period [88]. Taken together, our findings suggest a stronger contribution of myocardial inflammation and fibrosis to HCM phenotype in TnT mutants aligned with experimental observations [32, 89].

Although mouse models could help us to uncover HCM molecular mechanisms and treatments, differences in mouse and human heart functions can challenge the direct application of these findings to human patients [90, 91]. Moreover, mouse models usually show heightened cardiac hypertrophy than human HCM, and cardiac fibrosis, a key feature of human HCM, is typically less pronounced in mice [91]. Using the HCM model, we assessed the potential differences between mouse and human models of HCM and predicted molecular reactions that might be more pronounced in HCM patients, such as the regulation of Ca^2+^ sensitivity by titin, activation of IRS1 by growth factor receptors, and OGT regulation by PGC1*α*. Earlier studies indicate that titin-induced passive tension can elevate Ca^2+^ sensitivity, affecting sarcomere mechanics [92]. Considering the elevation of titin stiffness in both mouse and human models of HCM [93], our results suggest that the repercussions of higher titin stiffness could be more severe in HCM patients compared to HCM mouse models. Growth factors activate the PI3K/AKT pathway via IRS1, promoting physiological growth, survival, and glucose uptake and utilization in cardiomyocytes[94, 95]. Thus, the predicted increase in IRS1 activation by growth factor receptors in HCM patients compared to HCM mouse models, especially MyHC mutants, could indicate an elevated compensatory response in HCM patients through higher physiological hypertrophy, cell survival, and glucose metabolism. This amplified compensatory response may partly explain why some HCM patients remain asymptomatic or only exhibit symptoms during physical activity [81]. Regarding the projected heterogeneity in O-GlcNAc transferase (OGT) regulation by PGC1*α*, Umapathi et al. [96] showed that transgenic mouse hearts with overexpressing OGT exhibit higher O-GlcNAcylation leading to severe dilated cardiomyopathy, ventricular arrhythmia, and sudden death. In contrast, OGT overexpression could promote cell survival and mitigate oxidative stress and calcium overload following ischemia/reperfusion [97, 98]. Therefore, further research is needed to evaluate the impacts of a potential increase in OGT regulation by PGC1*α* in HCM patients.

### 3.4 Limitations and future directions

The computational systems biology model of HCM effectively predicted signaling variations, metabolic changes, and transcriptomic data consistent with experiments. It projected the shift from fatty acids to carbohydrates in HCM metabolism and highlighted the effects of HCM-induced ATP synthase deficiency on cardiomyocyte response. The model also shed light on mechanisms behind cardiomyocyte hypertrophic growth and apoptosis, identified shared and context-specific regulatory reactions that may control HCM phenotype across mutations and species, and suggested potential therapeutic targets for HCM. While insightful, the HCM model has some methodological limitations impacting its predictive abilities and its application to clinical data. Due to a lack of detailed kinetic data for familial hypertrophic cardiomyopathy signaling pathways, the model primarily relies on default parameters and captures steady-state responses of signaling entities. This means it may not depict the evolution of signaling changes or distinguish primary from secondary signals accurately. However, the model still shows strong validation in the steady-state responses crucial for chronic diseases like HCM (refer to Fig. 2). Moreover, building the model mainly on causal interactions between cardiomyocyte signaling entities and using default parameters inherently addresses the prevalent problem of overfitting to a training dataset. Additionally, while a dynamic cardiac metabolic network model is ideal, its creation is challenging because of the large scale of this network and scarce kinetic data. Consequently, the integrated iCardio metabolic network model might not capture metabolic pathway dynamics or reflect metabolite level changes.

In future studies, integration of cardiomyocyte and sarcomere mechanics models with the current HCM model could offer more insights into the HCM genotype-to-phenotype mechanisms, especially for HCM mutations whose primary effects are other than increased Ca^2+^ sensitivity. Furthermore, considering the critical role of fibrosis in the HCM phenotype, a multiscale model capturing processes from the molecular dimension (e.g., signaling pathways leading to fibroblast activation and collagen synthesis) to the tissue and organ scales (e.g., changes in cardiac tissue stiffness and heart function) could yield a comprehensive understanding of cardiac fibrosis in HCM. This multiscale model would integrate components such as cardiomyocytes, fibroblasts, the extracellular matrix, and intercellular communications to predict cardiac fibrosis in the context of HCM. Finally, developing a multiscale spatiotemporal HCM model that encompasses the cardiac remodeling dynamics from the molecular to organ scales and yields clinical indicators like ejection fraction could be a valuable tool for HCM patient-focused clinical research.

## ACKNOWLEDGMENTS

The authors thank Dr. Jason Papin for his valuable input in shaping the concept and design of this research and Drs. Emmet Francis and Kim McCabe for their feedback and thorough review of this paper. This work was supported by an AHA postdoctoral fellowship (ID: 898850) to A.K., and in part by the Wu Tsai Human Performance Alliance at UCSD to P.R.

## AUTHOR CONTRIBUTIONS

A.K.: Conception and design of the work; Acquisition, analysis, and interpretation of data for the work; Drafting the work; Final approval of the version to be published; Agreement to be accountable for all aspects of the work. J.J: Conception and design of the work; Review and edit of the manuscript; Final approval of the version to be published; Agreement to be accountable for all aspects of the work. P.R.: Conception and design of the work; Revising the work critically for important intellectual content; Review and edit of the manuscript; Final approval of the version to be published; Agreement to be accountable for all aspects of the work.

## STAR METHODS

### RESOURCE AVAILABILITY

#### Lead contact

Further information and requests for resources and reagents should be directed to and will be fulfilled by the corresponding author, Padmini Rangamani.

#### Materials availability

This study did not generate new unique reagents.

#### Data and code availability

The code generated during this study is available at https://github.com/mkm1712/HCM-Model. The code used for sensitivity analysis is available at https://github.com/mkm1712/Morris-Sensitivity-Analysis. The code used for simulating the metabolic network model (iCardio) is available at https://github.com/csbl/iCardio. The Netflux software is available at https://github.com/saucerman-lab/Netflux.

### METHOD DETAILS

#### Development of model structure

This section describes the main features of the HCM systems biology model. The HCM model was developed by modeling integral systems within the cardiomyocyte that jointly modulate genotype to phenotype mechanisms in hypertrophic cardiomyopathy. These systems include 1) the signaling network, 2) the metabolic network, 3) the gene regulatory network, and 4) the post-translational crosstalk between signaling and metabolic networks. For the cardiomyocyte signaling, we expanded the familial cardiomyopathy signaling network model previously developed by the authors [24] by adding mediators that link the cardiomyocyte signaling to its metabolic and gene regulatory networks. These mediators and their interactions with the original signaling network model [24] were manually curated from the literature (See Supplementary Table 3).

For the metabolic network model, we adopted a heart-specific genome-scale metabolic network model (iCardio) [10] that determines significantly altered metabolic functions based on cardiomyocyte transcriptome using the TIDEs (Tasks Inferred from Differential Expression) approach, for more details on the iCardio model and the TIDEs approach refer to [10]. However, briefly, the TIDEs approach as a reaction-centered methodology employs metabolic tasks and their associated reactions, identified using iCardio with pFBA (parsimonious Flux Balance Analysis), to determine metabolic functions that exhibit significant associations with differentially expressed genes in a specific condition. The network reactions are weighted by overlaying gene expression log fold changes onto them, employing GPR rules that represent the proteins required for catalyzing specific reactions through AND or OR relationships. The task scores are calculated by averaging assigned reaction weight values within a task. To determine the statistical significance of task scores, gene expression fold changes are randomized 1000 times among the genes in each dataset, and task scores are recalculated using the randomized data, creating a distribution of task scores. The significance (p-value) of each task score is determined by comparing the number of random task scores greater/less than the original data, depending on the position of the task score relative to the mean randomized task score. A p-value < 0.05 was used for determining the significance of task scores [10].

To link expanded signaling and metabolic network models through a transcriptional regulatory network model, we first methodically reviewed the literature to identify major transcription factors that regulate the level of mRNAs associated with metabolic enzymes in the iCardio model as well as transcription factors regulating cardiomyocyte hypertrophic growth and remodeling [44, 99–101]. Then, we utilized databases of transcriptional regulatory networks (i.e., TRRUST [102]) and binding motif analysis tools (i.e., iRegulon [103]) to identify candidate target genes directly regulated by these transcription factors. Transcriptional regulation has been confirmed by published co-expression experiments and analyses of ChIP-seq datasets in mice or rates. The expression of genes at the transcriptional regulatory network model was used as the input for the iCardio model, considering the assumption of linear association between mRNA and protein levels for metabolic enzymes in the iCardio model. However, for metabolic enzymes regulated by both changes in the mRNA level and post-translational modifications in the model, a simple formula was used to combine these effects (i.e., *log*(*FC_A_*) = *log*(*FC_mRNA_*) + *log*(*FC_post_*)). We also included the feedback from mRNAs to signaling species in the model.

In summary, the systems biology model of HCM, excluding the iCardio model, comprises 44 signaling species, 15 metabolic mediators, 23 transcription factors, 1078 genes, and 1307 reactions, including 40 feedback reactions from genes to their associated proteins. The iCardio heart metabolic network model includes 4200 reactions, 2890 metabolites, and 1734 genes predicting changes in 347 metabolic tasks/functions[10]. In addition to the HCM mutation input, the HCM model also includes two environmental inputs (i.e., pressure overload and volume overload) to evaluate cardiomyocyte response in different contexts.

#### Mathematical model of HCM

We used a stochastic differential equation approach to model the expanded signaling and gene regulatory networks. For the deterministic part, we used the logic-based differential equation (LDE) approach [104] to convert the interaction between species to a system of ordinary differential equations. The LDE approach was frequently used to model large-scale signaling networks with limited available data [11, 24, 104–107]. In this approach, we used Logical AND (i.e., *f* (*x*)*.f* (*y*)) or OR operations (i.e., *f* (*x*) + *f* (*y*) *− f* (*x*)*.f* (*y*)) to model pathways crosstalk. The AND gate is used when each input to a node is necessary for its activation, whereas the OR gate is used when each input is sufficient but not necessary for the node activation. The default parameters of Hill coefficient n = 1.4 and half-maximal effective concentration EC50 = 0.5 were used for all reactions. Default node parameters included initial activation Yinit = 0.01 and maximal activation Ymax equals 1 for signaling species and 0.5 for mRNAs. To account for the distinct time scales of signaling species, mRNAs, and phenotypic changes like cell area, we employed three default time constants, *τ* = 1 for signaling species, *τ* = 10 for mRNAs, and *τ* = 20 for cell area. Based on data availability, these time constants can be estimated to reproduce similar time scales of species observed in experiments. Considering the substantial size of the model, we utilized four default weights (*W* = 0.1; 0.2; 0.5; 1) based on the type of reaction. This approach was adopted to maintain a balance between the responsiveness of the model to inputs (low weight decreases model responsiveness) and its potential for becoming saturated (high weights saturate activity of model species). For example, in reactions with the AND logic, we used *W* = 1 to compensate for the decrease in the signal strength after an AND gate. The Netflux software has been used to generate the system of LDEs based on a predefined Microsoft Excel format defining model species, reactions, and parameters (Supplementary Table 3).

To account for the extrinsic biological noise in the signaling and gene regulatory networks, a noise term was added to the differential equations of each species using the Ornstein-Uhlenbeck process [108] (see Eq.1).

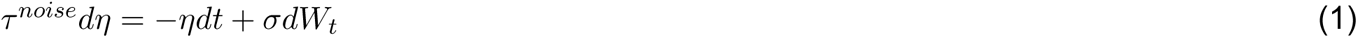

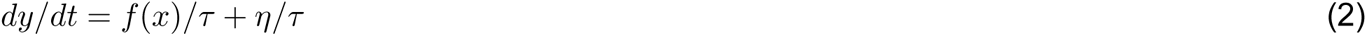

Where *τ^noise^*and *σ* are noise timescale and amplitude, respectively. *W_t_* are identically distributed Weiner processes independent for each species [109].*η*(*t*) has a zero mean and *σ*^2^*/*2*τ^noise^*variance. *τ^noise^* was selected equal to one for the model simulations. The noise term was then numerically solved using the Milstein scheme [110]. To solve the dynamics of the nodes (see Eq. 2) over the simulation time interval, the Euler scheme was employed. Given that the noise term has a zero mean, for reproducibility, we utilized deterministic fold changes calculated by LDE for analyses of mRNAs, except for the volcano plot (Fig. 3B) where the mean of *in silico* replicates was used.

In order to establish the basal condition (control state) of the cardiomyocytes, we ran the model without HCM input until we achieved a steady state. Then, HCM input was applied to simulate hypertrophic cardiomyopathy. Fifty (50) *in silico* replicates were simulated per condition with a noise amplitude equal to 0.05 for signaling species and 0.25 for mRNAs. We used higher noise amplitude for genes to compensate for the effect of their longer timescales on noise magnitude (see Eq. 2) compared to signaling species. The median coefficient of variation for mRNAs was calculated as 8%, aligning with the range observed in experimental data [111]. The inverse relationship between the sensitivity of model species to the input and the number of mediators meant that the simulated alterations in mRNA levels were significantly lesser compared to signaling species upstream. To compensate for this reduced sensitivity, we amplified changes in mRNAs by increasing their Ymax by using a power function (see Eq. 3). Selecting *β* = 2 led to changes in mRNA levels that were within the physiological range and comparable to those changes seen in the signaling model species. Notably, this amplification does not alter the direction or significance of gene expression. All simulations were conducted using MATLAB 2021.

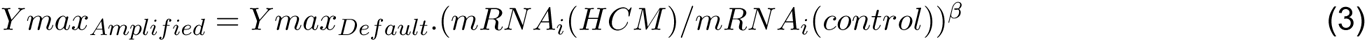

#### Development of hypertrophic growth and apoptosis indexes

To quantitatively analyze cardiomyocyte remodeling within the context of HCM, we developed two phenotypic indexes based on variations in regulatory mRNA expressions within the *in silico* transcriptome. These expressions regulate hypertrophic growth and apoptosis in cardiomyocytes. The mRNA expressions were identified from gene sets (Gene Ontology or GO) correlated with either promoting or inhibiting cardiac muscle hypertrophy (i.e., GO:0010613, GO:0061051, GO:0014898, GO:0003300, GO:0061049, GO:1903243, GO:0061052, GO:0010614, GO:0014898), hypertrophy markers/regulators [112, 113], and gene sets associated with the regulation of apoptosis in cardiac muscle cells (GO:0010665). The expression levels of these mRNAs, depending on their promotive or inhibitive role in managing cardiomyocyte hypertrophic growth or apoptosis, either stimulated or repressed the corresponding indices within the model via hypothetical reactions assigned with a weight factor of 0.1. Given the significance of Nppa, Nppb, and Myh7 mRNAs as markers of a hypertrophic phenotype, their expression led to an activation of the hypertrophic growth index with a twofold weighting factor (w=0.2), as shown in the supplementary Table 3.

#### Morris global sensitivity analysis

We employed the Morris global sensitivity analysis approach to identify major reactions regulating cardiomyocyte response in each studied context. The Morris elementary effects method is a statistical strategy for screening large-scale models with many parameters or high computational requirements for model simulation. Employing the Morris method allows us to ascertain whether a parameter’s modification imparts a negligible or significant impact on the model’s output and whether this influence is linear or non-linear. Additionally, it enables us to identify whether the impact of a parameter is autonomous or interdependent with other parameters [114].To generate the sampling data, we applied the Sampling for Uniformity (SU) methodology with an input factor level of eight, an oversampling size of 300, and a trajectories number of 16, using a previously developed EE sensitivity package [43]. We chose the reaction weight (W) as the variable parameter for computing the Morris index (μ*) and standard deviation (σ) for each reaction. The parameters μ* and σ represent the mean of the absolute values and standard deviation of the elementary effects (EEs) [43]. The Morris index (μ*) quantifies the magnitude of each reaction’s influence on the chosen model output, while the standard deviation (σ) encapsulates its interdependence with other reactions, indicating non-linearity.

#### Enrichment analysis

Gene set enrichment analysis evaluates the collective behaviors of genes in terms of health and diseases [115]. Gene ontologies (GO) of differentially regulated genes in *in silico* transcriptome were retrieved through Enrichr (https://maayanlab.cloud/Enrichr/). Enrichr is a simple web-based enrichment analysis platform that provides various collective functions of gene lists [36]. GO Biological Processes 2023 and ClinVar 2019 databases were selected for GO analysis.

#### Statistical analysis

After data preparation (removing potential Inf or Nan) and imputation, we conducted an unpaired t-test between Control and HCM conditions to determine differentially expressed genes from *in silico* cell replicates. Calculated p-value (two-tailed) for each gene was employed to compute the false discovery rate (FDR) [116] in MATLAB 2021. Genes that exhibited an FDR of less than 0.05 were considered to be differentially expressed. The same approach was used to determine significant changes in the activity of signaling species. For statistical analysis with multiple conditions (refer to Fig. 4 and Fig. 6), we utilized ordinary one-way ANOVA followed by either Dunnett’s or Tukey’s tests for multiple comparisons with single pooled variance in GraphPad Prism v10.

